# *In silico* analysis of the fast-growing thermophile *Geobacillus* sp. LC300 using a novel genome-scale metabolic model

**DOI:** 10.1101/2021.04.07.438430

**Authors:** Emil Ljungqvist, Martin Gustavsson

## Abstract

Thermophilic microorganisms show high potential for use as biorefinery cell factories. Their high growth temperatures provide fast conversion rates, lower risk of contaminations, and facilitated purification of volatile products. To date, only a few thermophilic species have been utilized for microbial production purposes, and the development of production strains is impeded by the lack of metabolic engineering tools. In this study, we constructed a genome-scale metabolic model, *iGEL601*, of *Geobacillus* sp. LC300, an important part of the metabolic engineering pipeline. The model contains 601 genes, 1240 reactions and 1305 metabolites, and the reaction reversibility is based on thermodynamics at the optimum growth temperature. Using flux sampling, the model shows high similarity to experimentally determined reaction fluxes with both glucose and xylose as sole carbon sources. Furthermore, the model predicts previously unidentified by-products, closing the gap in the carbon balance for both carbon sources. Finally, *iGEL601* was used to suggest metabolic engineering strategies to maximise production of five industrially relevant compounds. The suggested strategies have previously been experimentally verified in other microorganisms, and predicted production rates are on par with or higher than those previously achieved experimentally. The results highlight the biotechnological potential of LC300 and the application of *iGEL601* for use as a tool in the metabolic engineering workflow.

## 1. Introduction

Thermophilic microorganisms show great potential as hosts in biorefinery processes producing various biochemicals and proteins. Their high growth temperatures can yield process advantages in large-scale, including increased reaction rates, lowered cooling costs, decreased risk of contaminations and facilitated purification of volatile products [1]. *Geobacillus* is a genus of aerobic, thermophilic, Gram-positive bacteria with typical optimal growth temperatures ranging between 55 – 60°C, and in some cases up to 75°C [2]. Due to their capabilities of consuming a diverse range of substrates including hexoses, pentoses, sugar polymers and fatty acids [2, 3], *Geobacilli* show potential as microbial biorefinery hosts. Despite this potential, only a few examples of metabolic engineering of *Geobacillus* for the production of industrial biochemicals have been reported [4, 5, 6, 7, 8]. The lack of metabolic engineering efforts could be explained by the limited availability of engineering tools for *Geobacillus* and thermophilic organisms in general [1]. Several recent publications report an expansion of this engineering toolbox, such as improving expression plasmids, discovery and adaptation of native *Geobacillus* CRISPR systems [9, 10] and synthetic promoter and ribosomal binding site libraries for tuning expression and translation [11]. In light of the increased availability of metabolic engineering tools, the prospect of implementing *Geobacilli* in biorefineries draws closer.

Genome-scale metabolic models (GEMs) facilitate prediction of a cell’s metabolic fluxes and can as such be used for prediction of different metabolic scenarios. These could be the effect of a certain drug on a human cell or the effect of a gene knock-out in a bacterium. Thus, GEMs can be powerful tools when combined with metabolic engineering efforts to guide the researcher to, for example, the optimal gene target to knock out for an optimized production of a certain product [12]. To create a GEM, the proposed genes of an annotated genome are assigned a function based on their homology to a database of potential orthologs. Compared to mesophilic organisms, thermophiles often have small genomes and amino acid sequences are usually short [13]. Furthermore, thermophilic proteins tend to differ in amino acid composition from their mesophilic counterparts, for example favoring charged residues over polar ones to increase stability through salt bridges [14, 15]. This difference in amino acid chain length and composition complicates homology-based metabolic modeling of thermophilic organisms, since this relies on using similarity of the protein sequence to assign it to a group of orthologs to predict its function. Since the majority of characterised enzymes stem from mesophilic organisms, assigning thermophilic protein sequences can thus be problematic. Despite these difficulties several GEMs of thermophilic organisms exist, including *Clostridium thermocellum* [16, 17], *Parageobacillus thermoglucosidasius* [18] and *Geobacillus iciganus* [19].

*Geobacillus* sp. LC300 (hereafter LC300) is a recently discovered thermophile showing promise for biorefining. LC300 displays fast growth (doubling times under 30 minutes) on both xylose and glucose, the two main sugars of lignocellulosic biomass, in defined media [20]. Due to these high substrate utilization rates, LC300 poses an interesting subject for metabolic engineering to rapidly convert substrates to renewable products of interest [20]. Although a central carbon metabolism model of LC300 has previously been published [20], and flux analysis of the central metabolism has been performed [21, 22], further insight in the central metabolism is required. For example, the core metabolism models overestimate biomass formation rates by 30%, and analysis of data from these previous reports reveal that this results from that the carbon and degree of reduction balances are still to be closed (Supplementary Table 6 & 7). Such metabolic knowledge is key to guiding rational metabolic engineering approaches, and genome-scale metabolic models are important tools in such endeavors [23].

To this end, we present a homology-based genome scale metabolic model of LC300. The model has reaction reversibilities set using thermodynamics data at LC300’s optimal growth temperature (72°C), and predictions validated against experimental data. Flux sampling of the model provided insight to further close the carbon and degree of reduction balances. Finally, flux sampling was used to evaluate the potential of LC300 as a biorefinery host and propose metabolic engineering strategies to optimise production of five industrially relevant biochemicals.

## 2. Materials & Methods

### 2.1. Model generation

A draft model of LC300 was generated using the RAVEN (Reconstruction, Analysis and Visualization of Metabolic Networks) toolbox 2 (v. 2.3.1) [24] for MATLAB [25], which constructs a model based on homology of protein sequences to orthologs in the KEGG (Kyoto Encyclopedia of Genes and Genomes) [26] database. The predicted protein coding sequences of LC300 (see GitHub repository) were downloaded from the NCBI genome database [27] (NCBI Accession number: ASM119162v1). The sequences were used as input in RAVEN, which parsed them against KEGG, assigned each compatible sequence to an ortholog in KEGG, and constructed a draft model.

### 2.2. Model development

The automatically generated model contained several gaps preventing synthesis of necessary biomass precursors. Since LC300 growths in defined media, it seemed unlikely that such metabolic gaps would exists. To verify that these genes indeed exits in LC300, the amino acid sequences of the corresponding genes from the close relative *Geobacillus stearothermophilus* were manually aligned against LC300 using NCBI BLAST [28], and all either resulted in hits with an E-value of less than 10^−20^ or were found to be already annotated in the LC300 genome (Supplementary Table 2). Thus, these genes and corresponding reactions were manually added to the model. Furthermore, exchange reactions and import reactions (e.g. ABC importers) of medium metabolites were added manually, along with the electron transport chain (ETC) and ATP synthase. The ETC reactions were constructed as follows:

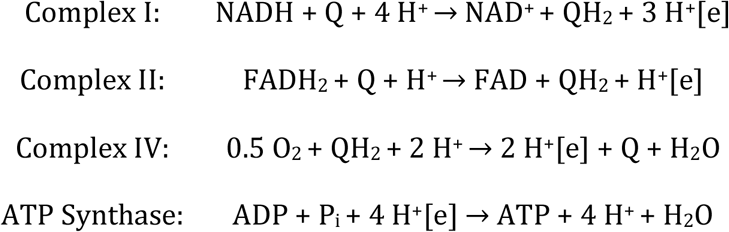

These reactions resulted in a predicted P/O-ratio of 1.25 for the oxidation of NADH and 0.75 for the oxidation of FADH_2_. The automatically generated model contained several generic metabolites without a determinable molecular mass (e.g. Acyl-CoA). Such metabolites were removed, along with the reactions they took part in.

The metadata of each metabolite was collected from MetaCyc [29], since the KEGG database lacks information about metabolite charge and uses metabolite formulae of fully protonated metabolites. For metabolites not found in the MetaCyc database (145 metabolites), a script developed by Chan *et al.* [30] was used to calculate metabolite formulas and charges automatically. This script was also used to confirm the molecular weight of the biomass component to 1 g/mmol, which results in the flux of the biomass reaction correctly corresponding the the specific growth rate *μ* (h^-1^). One issue in creating models of thermophiles is that reaction reversibilities in common databases are typically annotated for mesophilic growth temperatures. Thus, these are not necessarily valid at the optimum growth temperature of LC300. To alleviate this issue, the reversibility of reactions was determined based on thermodynamic calculations using the eQuilibrator API [31], using a script developed by Kumelj *et al*. [32]. The reaction conditions were set to pH 7, ionic strength 0.25 M and temperature 345 K. To evaluate reversibility, the oxidation of malate to oxaloacetate was used as a reference. This reaction has a positive ΔG’^m^ of 30 kJ/mol but is still driven in the forward direction by differences in metabolite concentration [33]. Since intracellular metabolite concentrations are unknown, this value of ±30 kJ/mol was selected as a cut-off for reversibility of each reaction. Using eQuilibrator, ΔG’^m^ predictions were obtained for 1074 of the 1155 non-exchange reactions in the model (Supplementary Table 4). This reversibility prediction was incorrect for some reactions (Supplementary Table 5), which had to be corrected manually. The remaining reactions not predicted by eQuilibrator were left at their default reversibility as annotated in the KEGG database.

### 2.3. Biomass objective function

The biomass objective function (BOF) was mainly based on data from the ^13^C-analysis of the LC300 central metabolism performed by Cordova *et al*. [22]. The BOF was constructed as a lumped reaction of protein, RNA, DNA, cell membrane, cell wall, glycogen and metabolic cofactors, along with the estimated necessary growth-associated ATP cost. The protein part of the BOF was created by all 20 amino acids along with the hydrolyzation of 1 mol ATP to 1 mol AMP and 2 mol GTP to 2 mol GDP per mol of amino acid. The respective concentration of each amino acid was set based on the data from Cordova *et al.* [22], with the addition of Arg, Cys, His and Trp which were excluded in the analysis of the beforementioned paper. The total amount of these amino acids was set to <1% of the total amount of amino acids, to deviate from the measured amino acid composition as little as possible. The RNA and DNA fractions of the BOF were constructed based on the biomass composition measured by Cordova *et al*., with RNA constructed of XMPs and DNA of dXMPs. RNA and DNA compositions were based on the GC-content of the LC300 genome. The cell membrane part of the BOF was constructed from fatty acids (tetradecanoic acid, palmitic acid and octadecanoic acid) and 0.5 mol glycerol phosphate per mol of fatty acids. The cell wall part of the BOF was constructed of 45% (w/w) peptidoglycan and 55% teichuronic acid. Since the LC300 genome lacks an annotated glycerol-3-phosphate cytidylyltransferase, an enzyme necessary for teichoic acid synthesis, teichuronic acid was chosen as the major cell wall component instead. In turn, peptidoglycan was constructed of N-acetylmuramoyl-(N-acetylglucosamine)-L-alanyl-D-glutamyl-meso-2,6-diaminopimeloyl-Dalanyl-D-alanine, and teichuronic acid of N-acetyl-D-galactoseamine and D-glucoronate. The components of the cofactor pool (Supplementary Table 1) were determined based on the work of Xavier *et al*. [34]. The concentrations of the components were based on previous work by Oh *et al*. [35]. The final model was named *iGEL601*.

**Table 1:** Comparison of predicted specific productivities by *iGEL601* growing at 75% of the maximum growth rate and highest reported specific productivities in literature, using glucose or xylose as carbon source.

| Specific productivity (mmol/gDW/h) |  |  |  |  |
| --- | --- | --- | --- | --- |
| Product | Carbon source | *iGEL601* predicted | Reported in literature | Reference |
| 3HB | Glucose | 15.7 | 1.1 | Perez *et al.* [39] |
|  | Xylose | 11.8 | 0.12* | Guaman *et al.* [43] |
| Ethanol | Glucose | 25.9 | 59.8* | Ruchala *et al.* [40] |
|  | Xylose | 18.6 | 10.7* | Qureshi *et al.* [41] |
| Fumarate | Glucose | 23.2 | 2.1* | Swart *et al.* [44] |
|  | Xylose | 16.5 | 0.7* | Wen *et al.* [45] |
| Lactate | Glucose | 25.9 | 9.6* | Bai *et al.* [42] |
|  | Xylose | 18.6 | 11.4* |  |
| Succinate | Glucose | 19.8 | 22* | Lee *et al.* [46] |
|  | Xylose | 15.2 | 1.6* | Olajuyin *et al.* [47] |

\*Calculated from data presented in reference

### 2.4. Calculation of balances

Calculations on carbon balance and degree of reduction[36] was performed on the data published by Cordova *et al*. [21, 22], where the degree of reduction is a measure of the electron-donating capabilities of each compound. Each compound is assigned a value based on its chemical formula (represented by *γ*), and the sum of the degree of reduction of the in-going compounds must match the sum of the outgoing compounds in any system. For example, the degree of reduction of carbon dioxide is calculated in equation 1. Calculations of degree of reduction and elemental balances for LC300 growing on glucose and xylose are shown in Supplementary Tables 6 and 7, respectively.

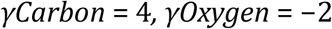

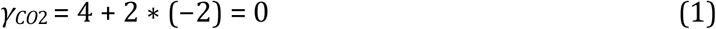

### 2.5. Flux Balance Analysis

Parsimonious flux balance analysis (pFBA) attempts to solve the equation system that is the GEM, while simultaneously optimizing the objective function and minimising the sum of all fluxes. pFBA was performed on *iGEL601* using the RAVEN toolbox 2.0 [24]. Two pFBA runs were performed, using glucose and xylose as carbon sources respectively. For both runs, biomass formation was set as the objective function.

### 2.6. Flux sampling

Flux sampling was performed using the ACHR sampler in the COBRA toolbox 3.0 [37]. Prior to executing the sampling algorithm, the maximum biomass formation rate was determined using pFBA, and all blocked reactions in the pFBA solution were removed so simplify the sampled model. The sampling options were set to: 200 steps per point, 1000 points returned, 5000 warmup points.

### 2.7. Modifying iGEL601 for production of biochemicals

All but one of the reactions necessary for (R)-3-hydroxybutyrate (3HB) synthesis were found in *iGEL601*. The last reaction, hydrolysation of (R)-3-hydroxybutyryl-CoA to the free fatty acid, is usually performed by medium-chain acyl-CoA thioesterases. Three such thioesterases are annotated in the LC300 genome (IB49 00680, IB49 01350, IB49 18150) and the 3HB thioesterase reaction was assigned to the genes of all three. Furthermore, exchange reactions of 3HB, ethanol, fumarate, and succinate were added to allow net production of these metabolites. The resulting production models were sampled as described above.

## 3. Results

### 3.1. Model generation

A draft model of LC300 was generated automatically based on protein homology. In this draft model 20 non-spontaneous reactions were found to be missing (Supplementary Table 2), leading to gaps preventing synthesis of necessary biomass precursor metabolites. The amino acid sequences of the *G. stearothermophilus* genes responsible for these reactions were manually aligned against the LC300 genome. All of these genes either resulted in hits with an E-value of less than 10^-20^ or were found to be already annotated in the LC300 genome. Based on this, it was considered likely that LC300 contained the genes necessary for these reactions, which is further supported by previous reports of growth in defined media [20]. Thus, the 20 missing reactions were manually added to the model. In line with earlier findings [20, 19], no gene for 6phosphogluconolactonase, catalysing a necessary reaction in the oxidative branch of the pentose phosphate pathway, was identified in the LC300 genome. Nevertheless, its catalysed reaction has been shown to be spontaneous at room temperature *in vitro* [2], and it was therefore included in the model.

After establishing the initial reaction network from the LC300 genome, exchange reactions and import reactions (Supplementary Table 3) of medium metabolites were added manually to allow for simulating the uptake of nutrients necessary for growth. These include exchange and import reactions of the four experimentally validated carbon sources (glucose, xylose, galactose and mannose). Furthermore, reactions for the electron transport chain (ETC) and ATP synthase were manually added to the model, with the composition of the ETC detailed in the Materials and Methods section. Finally, the biomass objective function was constructed as a lumped reaction of macromolecules consisting of protein, RNA, DNA, cell membrane, cell wall and glycogen, as well as necessary cofactors and ATP. The amount of each component in the biomass function was based primarily on the biomass composition reported previously [20], with additional assumptions detailed in the Materials and Methods (Figure 1A). The resulting elemental composition of the biomass component was verified to a molecular weight of 1 g/mmol, ensuring that the flux of the biomass reaction correctly corresponds to the specific growth rate *μ* (h^-1^). The final model, *iGEL601*, contains 601 genes, 1240 reactions and 1305 metabolites. The reaction distribution of *iGEL601* can be seen in Figure 1B.

**Figure 1:**
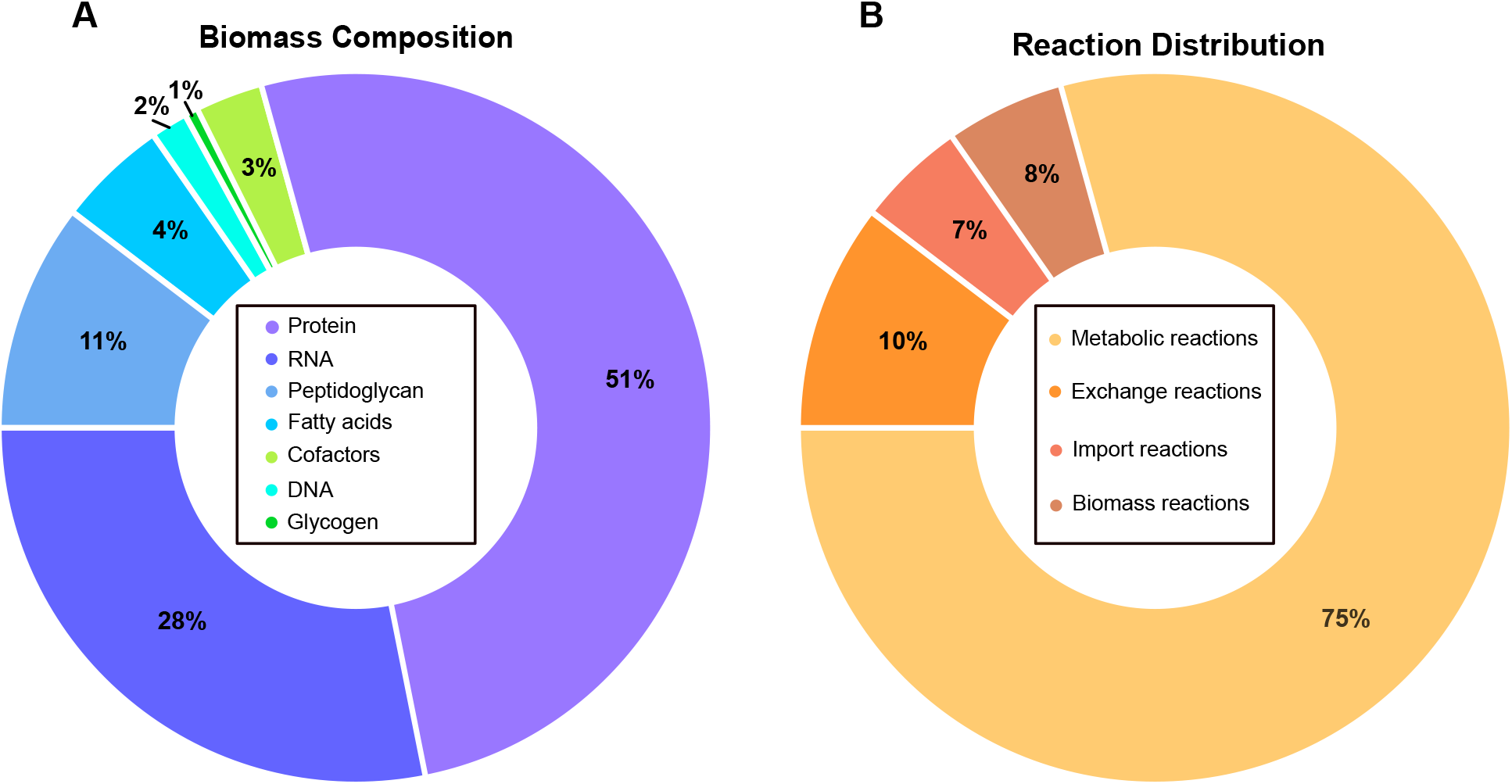
(A) Biomass composition of iGEL601. (B) Reaction distribution of iGEL601.

With the reactions and metabolites of iGEL601 established, the next step was to differentiate between reversible and irreversible reactions. This was performed based on thermodynamic calculations using the optimal growth temperature of LC300 (72°C) as input temperature. As a result, the reversibilities for 1074 of the 1155 non-exchange reactions in the model could be corrected. For the remaining 81 reactions, the default reversibilities obtained from KEGG were assumed. As a final step of the model generation, the model was fine-tuned to give accurate predictions of the growth rate of LC300. To this end, parsimonious flux balance analysis (pFBA) was performed on the model while constraining sugar uptake, oxygen uptake and acetate production to experimentally determined rates [21, 22], with biomass formation set as the objective function. Here it was verified that the model supports growth on all of the four experimentally validated carbon sources (glucose, xylose, mannose and galactose). Of these, detailed growth data was available only for glucose and xylose. This data was used to adjust the P/O-ratio of the ETC and the non-growth associated maintenance (q_m_), until the model gave accurate estimations of biomass yield. As a result, the P/O ratio on NADH was set to 1.25 and the P/O ratio on FADH_2_ was set to 0.75. Finally, accurate biomass yields was achieved through different q_m_ values for glucose (10 mmol/gDW/h) and xylose (50 mmol/gDW/h). With these parameters, the biomass yield on glucose and xylose slightly overestimated (by 2% and 7%, respectively). As a final validation step, *iGEL601* was analysed with the MEMOTE tool [38], showing close to 100% charge and mass balance.

### 3.2. Comparison of model predictions to experimental flux data

After constructing *iGEL601* and fitting it to predict reported experimental biomass yields, we proceeded to validate the predicted metabolic fluxes by comparison with published C13 metabolic flux data [21, 22]. Initially, a crude comparison was made based on pFBA analysis with high similarities between the predicted and experimental values (data not shown). However, due to the underdetermined nature of the linear equation system comprising GEMs such as *iGEL601*, the flux solution resulting from pFBA is but one of an infinite number of flux solutions inside the solution space. Thus, flux sampling was performed on the *iGEL601* solution space in order to get a more representative estimate of the predicted fluxes.

This sampling was done using glucose and xylose as carbon sources with the minimum biomass formation rate constrained to above 95% of the maximum obtained by pFBA on the different carbon sources. The predicted central carbon metabolism reaction fluxes (see GitHub repository for data) were compared to the previously reported experimentally determined fluxes [21, 22], as visualised in Figure 2. Overall, the reactions have small standard deviations, indicating a highly constrained solution space under the simulated conditions. One exception to this is the phosphoenolpyruvate (PEP) to pyruvate (PYR) conversion, where three separate reactions (utilizing ADP, dADP and dGTP respectively as cofactors) can carry flux. The flux of these reactions and their standard deviation have been combined to correctly represent the total flux going from PEP to PYR, and the standard deviation is increased as a result. Similarly, part of the succinyl-CoA to succinate conversion is done via the lysine biosynthesis pathway, and this flux has been added to the flux of succinyl-CoA synthetase like above. When glucose is used as carbon source (Figure 2A) the fluxes show close to perfect fit. When xylose is used as carbon source (Figure 2B) the overall fit is again good, though the y-axis intercept is set at −2.15. This indicates an overall underestimation of the predicted fluxes throughout metabolism of *iGEL601* as compared to the experimentally determined values. In contrast to the experimental data, *iGEL601* predicts acetate and lactate as byproducts. As these fluxes are not included in the flux comparison of Figure 2B, it is likely the cause of the general flux underestimation.

**Figure 2:**
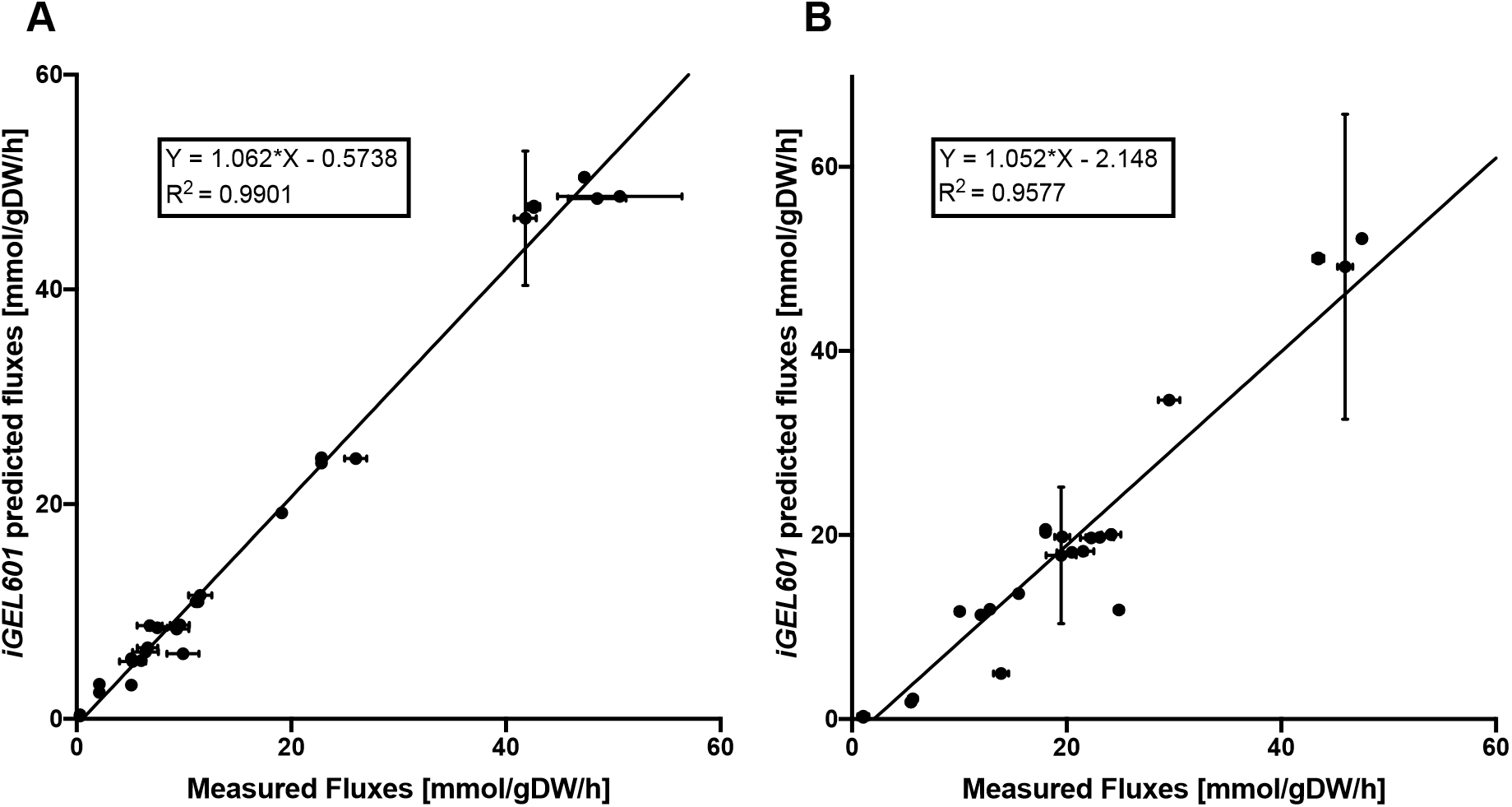
Comparison of estimated fluxes by Cordova *et al.* [21, 22] and predicted fluxes by flux sampling of *iGEL601*. (A) Flux sampling with glucose as carbon source, constraining glucose uptake, oxygen uptake and acetate production to experimentally determined values. (B) Flux sampling with xylose as carbon source, constraining xylose and oxygen uptake to experimentally determined values. Each point represents a reaction with its experimentally determined flux [21, 22] on the x-axis and the mean of the flux sampling of *iGEL601* on the y-axis. The standard deviation of each reaction is visualized as error bars. Linear regression with slope 1, intercept in origo and R^2^=1 corresponds to perfect fit.

### 3.3. Modelling the LC300 central metabolism

After validating *iGEL601* against the existing experimental knowledge, we proceeded to seek additional insight into the LC300 metabolism using the model. Previously reported flux data has left a gap in the carbon and degree of reduction balances, indicating the presence of hitherto unidentified byproducts of the LC300 metabolism. In order to identify potential by-product candidates, we enabled exchange reactions for export of all 20 amino acids, as well as a selection of central metabolites (acetate, lactate, pyruvate, citrate and succinate) and performed solution space sampling as described above. The resulting flux distribution in the central carbon metabolism can be seen in Figure 3. For both carbon sources, the common central carbon metabolism pathways are utilized (i.e. pentose phosphate pathway, glycolysis and citric acid cycle), with the main source of NADH being glyceraldehyde-3-phosphate dehydrogenase. The main source of NADPH is the oxidative pentose phosphate pathway when glucose is used as carbon source, while isocitrate dehydrogenase is the main source in the case of xylose.

**Figure 3:**
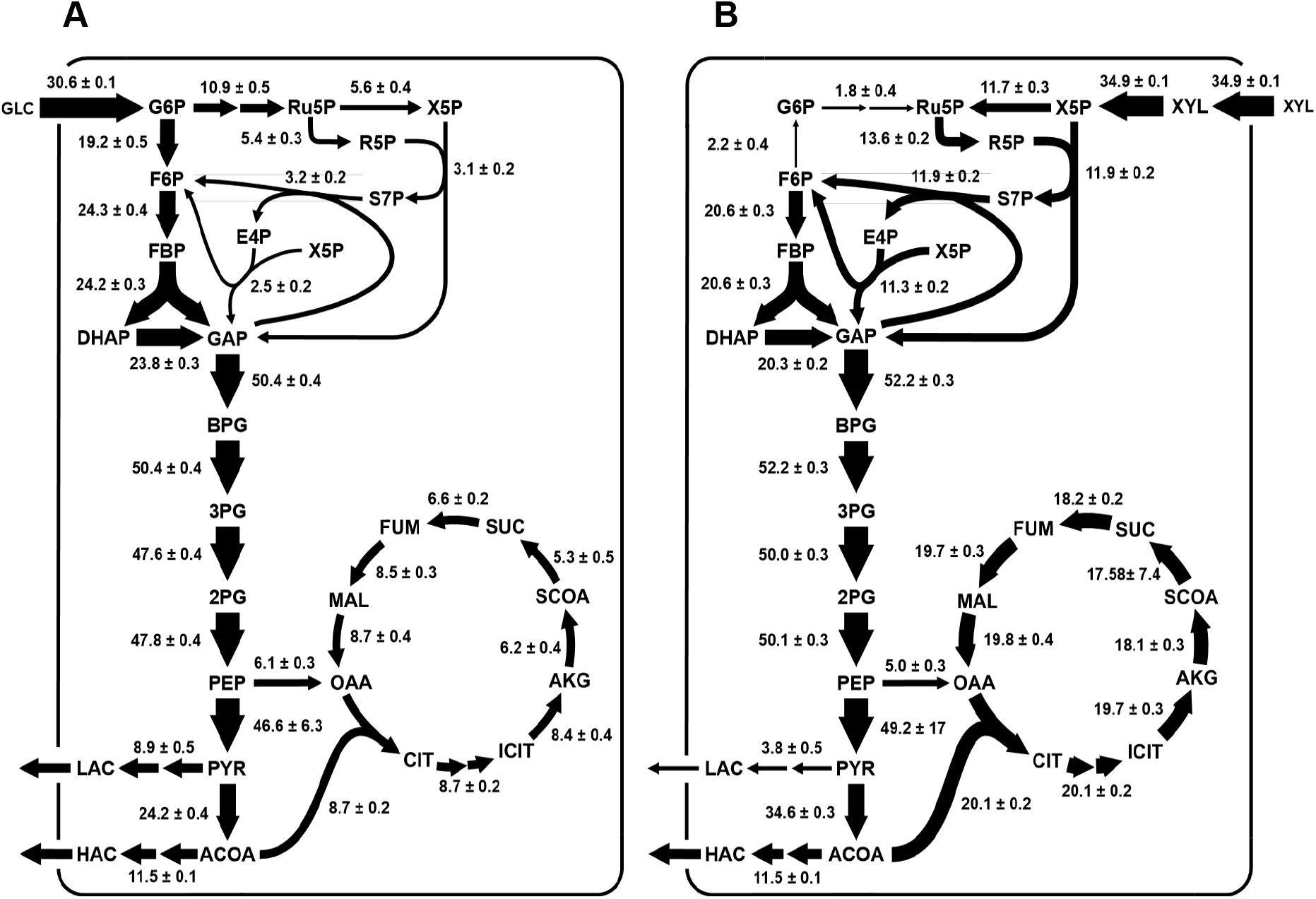
Flux sampling of iGEL601 with biomass as objective function. Displayed fluxes are means of the 1000 samples, plus/minus the standard deviation. (A) Glucose as carbon source, oxygen uptake and acetate production constrained to experimentally determined values. (B) Xylose as carbon source, oxygen uptake and acetate production constrained to experimentally determined values. Non growth-associated maintenance increased. *Abbreviations: G6P - Glucose-6-Phosphate, Ru5P - Ribulose-5-Phosphate, X5P - Xylulose-5-Phosphate, R5P - Ribose-5-Phosphate, S7P - Sedoheptulose-7-Phosphate, E4P Erythrose-4-Phosphate, F6P - Fructose-6-Phosphate, FBP - Fructose 1,6-bisphosphate, DHAP - Dihydroxyacetone phosphate, GAP - Glyceraldehyde-3-phosphate, BPG - 1,3-bisphosphoglycerate, 3PG - 3-phosphoglycerate, 2PG - 2-phosphoglycerate, PEP - Phosphoenolpyruvate, PYR - Pyruvate, ACOA - Acetyl Coenzyme A, LAC - Lactate, HAC - Acetic acid, CIT - Citrate, ICIT - Isocitrate, AKG - α-ketoglutarate, SCOA - Succinyl Coenzyme A, SUC - Succinate, FUM - Fumarate, MAL - Malate, OAA - Oxaloacetate*.

On both glucose and xylose, acetate and lactate are predicted byproducts. For glucose, the predicted production rates are 11.5 mmol/gDW/h acetate and 8.9 mmol/gDW/h lactate. The lactate production is in contrast with earlier reported experimental data [22], and could be responsible for a large part of the missing carbon in that data. On xylose the predicted rates are 11.5 mmol/gDW/h acetate and 3.8 mmol/gDW/h lactate. None of these byproducts are reported in earlier experimental data [21], but could also explain the large gaps in carbon and degree of reduction balances.) On both glucose and xylose, acetic acid is a predicted overflow metabolite (11.5 mmol/gDW/h and 10.9 mmol/gDW/h respectively. The addition of these byproducts in the balance calculations for carbon and degree of reduction, reduces the gaps in earlier reported data. In the case of glucose, the addition of lactic acid closes the carbon balance gap from 28% to 14% and the degree of reduction gap from 27% to 13% (Supplementary Table 6). In the case of xylose, the addition of acetic acid closes the carbon balance and degree of reduction balance gaps from 35% to 12% (Supplementary table 7). Here it should be noted that the additional active exchange reactions allowing export of the 20 amino acids, pyruvate, citrate and succinate also all carry minor fluxes (GitHub repository) under the simulated conditions. Together, these likely comprise the majority of the missing carbon and degree of reduction noted here.

### 3.4. Modeling for metabolic engineering

As *iGEL601* produced good estimations of experimental data, we finally attempted to assess the potential of the LC300 metabolism for production of biochemicals. Genome-scale models allow metabolic engineers to take a systems biology approach to design of strains [23], where the role of the model is to take genotypic information as input and converting it to phenotypic predictions as output. In this way, metabolic engineers can get hints of which genotypic alterations are needed to achieve a certain goal phenotype. At first glance LC300 has great potential for biorefinng due to its high substrate uptake rates of both hexoses and pentoses.

If some of this carbon flux could be diverted towards fuels or chemicals of industrial relevance through metabolic engineering, LC300 could potentially yield biorefinery strains with very high productivities.

To evaluate this potential *in silico*, *iGEL601* was modified to allow for production of 5 different industrially relevant biochemicals: (R)-3-hydroxybutyrate (3HB), ethanol, fumarate, lactate, and succinate. Flux sampling was performed with biomass formation locked at 75% of the maximum rate as determined by pFBA on glucose and xylose respectively. For each product of interest, flux sampling was performed with maximisation of the product exchange reaction set as objective. The fluxes of the central carbon metabolism were then compared to a reference sampling with biomass formation locked at 95% of optimum. The resulting flux changes (shown in Figure 4) can give an indication of which pathways need to be up- or downregulated in order to achieve 25% diversion of consumed carbon to product.

**Figure 4:**
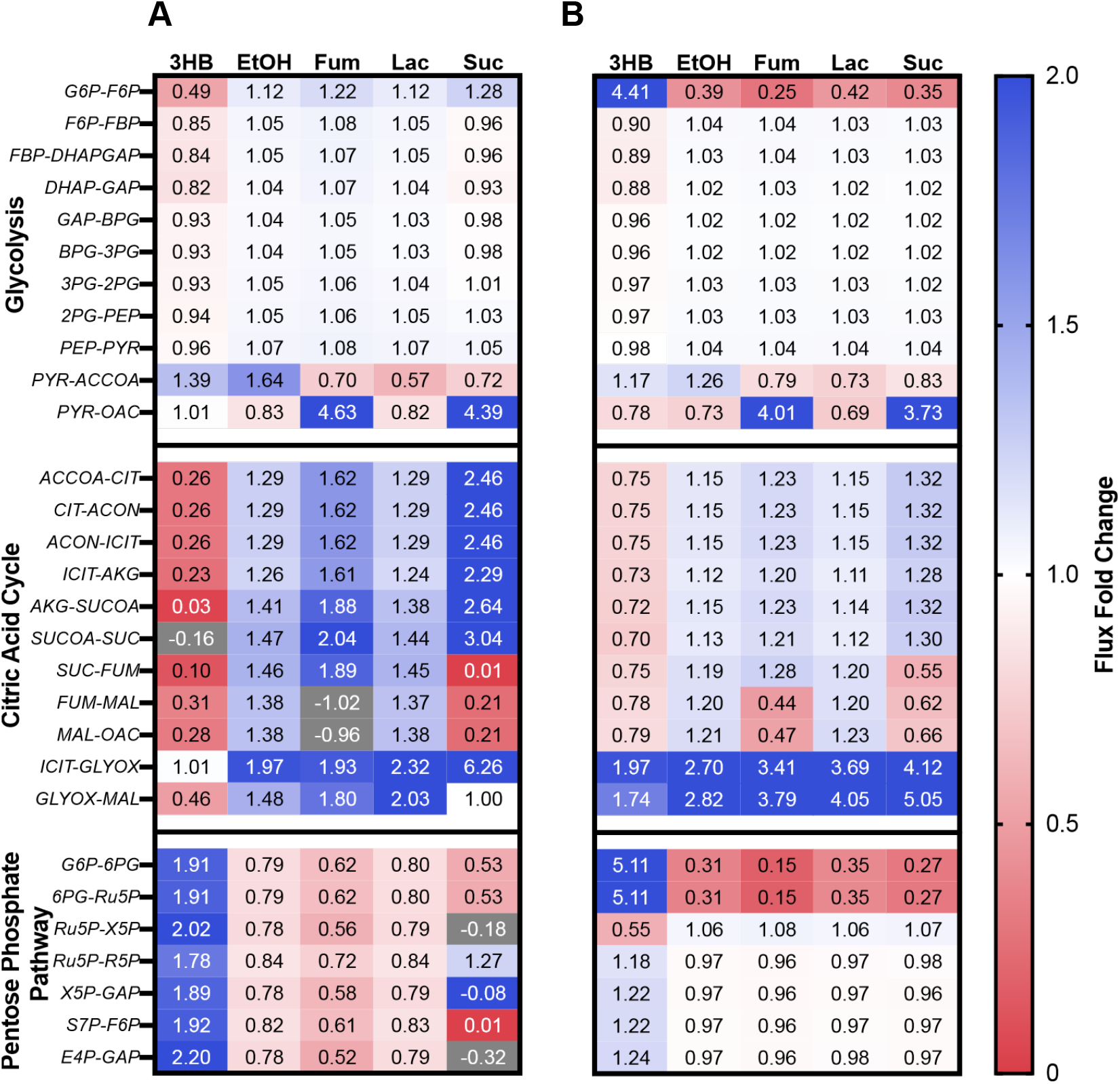
Flux fold change of reactions in central carbon metabolism, optimizing for production of the 5 different products. Gray color indicates negative flux changes i.e. reactions that have changed direction. (A) Glucose as carbon source. (B) Xylose as carbon source. *Abbreviations: 3HB - 3-Hydroxybutyrate, EtOH - Ethanol, G6P - Glucose-6-Phosphate, 6PG - 6-phosphogluconate, Ru5P - Ribulose-5-Phosphate, X5P - Xylulose-5-Phosphate, R5P - Ribose-5-Phosphate, S7P - Sedoheptulose-7-Phosphate, E4P - Erythrose-4-Phosphate, F6P - Fructose-6-Phosphate, FBP - Fructose 1,6-bisphosphate, DHAP - Dihydroxyacetone phosphate, GAP - Glyceraldehyde-3-phosphate, BPG - 1,3-bisphosphoglycerate, 3PG - 3-phosphoglycerate, 2PG – 2-phosphoglycerate, PEP - Phosphoenolpyruvate, PYR - Pyruvate, ACOA - Acetyl Coenzyme A, LAC - Lactate, HAC - Acetic acid, CIT - Citrate, ACON - Aconitate, ICIT - Isocitrate, AKG - α-ketoglutarate, SCOA - Succinyl Coenzyme A, SUC Succinate, FUM - Fumarate, MAL - Malate, OAA - Oxaloacetate. GLYOX - Glyoxylate*

In the case of 3HB production, the pentose phosphate pathway is upregulated on both sugars (Figure 4), which is likely a requirement in order to produce sufficient NADPH to drive the reductive pathway from acetyl-CoA to 3HB. Additionally, the flux through the citric acid cycle is downregulated in both cases, while the flux through glycolysis is maintained. With these modifications, along with deletion of acetic acid and lactic acid production, a 3HB production rate of 15.7 mmol/gDW/h is predicted on glucose and 11.8 mmol/gDW/h on xylose (Table 1). This is a 15-fold increase of the highest reported specific 3HB productivities of *Escherichia coli* [39] on glucose.

For ethanol and lactate production, glycolysis is slightly upregulated at the cost of flux through the pentose phosphate pathway, especially the oxidative pentose phosphate pathway on xylose. This is expected, as the lower modelled growth rate requires less NADPH for biomass production, leaving more sugar free to proceed through the glycolysis instead of the pentose phosphate pathway. The citric acid cycle is also moderately upregulated on both sugars. The predicted specific ethanol productivities on glucose and xylose were 25.9 mmol/gDW/h and 18.6 mmol/gDW/h, respectively (Table 1). For glucose, this is roughly half of the reported specific ethanol productivities in *Saccharomyces cerevisiae* [40], while on xylose, it is almost 2-fold of that reported in *Escherichia coli* [41]. The predicted specific lactate productivities on glucose and xylose were 25.9 mmol/gDW/h and 18.6 mmol/gDW/h, respectively. This is a more than 2-fold increase of the reported specific lactate productivity in *Lactobacillus lactis* [42].

For fumarate and succinate production, pyruvate carboxylase is heavily upregulated due to the requirement of replenishing oxaloacetate during during production of citric acid cycle-derived products. For fumarate production on glucose, the citric acid cycle is split into a forward, oxidative branch from citric acid to fumarate, and a reverse, reductive branch making fumarate from oxaloacetate. Together, this indicates that an anaerobic cultivation setting might be advantageous for this product, since the described behaviour of the citric acid cycle is common in fermentative metabolism. The predicted specific fumarate productivities on glucose and xylose were 23.2 mmol/gDW/h and 16.5 mmol/gDW/h, respectively (Table 1). This is 10-fold higher than reported specific fumarate productivities on glucose and 20-fold higher on xylose. For succinate, the predicted specific productivities were 19.8 mmol/gDW/h on glucose and 15.2 mmol/gDW/h on xylose (Table 1). This is slightly lower than the highest reported specific productivities in *Mannheimia succiniproducens* on glucose [46], but 10-fold higher than the reported specific productivities on xylose in *E. coli* [47].

For most cases, the glyoxylate shunt is additionally upregulated; however, since the flux through these reactions is minimal in the reference samplings, a 6-fold increase in the production samplings still results in fluxes close to zero.

## 4. Discussion

In this study we constructed a genome scale metabolic model, *iGEL601*, of *Geobacillus* LC300. The biomass composition was constructed based on experimental data, and the reaction kinetics were adapted to match the thermodynamics at the optimum growth temperature of the organism. The final model contains 601 genes, covering roughly 25% of the predicted protein coding part of the LC300 genome.

Previous studies have analyzed the flux distribution of LC300 using ^13^C MFA [21, 22]. When compared to these data, *iGEL601* shows great similarity throughout the central carbon metabolism (Figure 2) and respiration pathways of oxygen and carbon dioxide. Based on this, we conclude that the modelling assumptions made above seem reasonable, and consequently *iGEL601* could be used as a realistic estimate of the LC300 metabolism.

When glucose is used as carbon source, flux through the pentose phosphate pathway is especially high, possibly due to the high demand for NADPH in biosynthesis reactions. It is likely that a high flux through here is needed when growing at the high maximum growth rate of LC300. Acetic acid is known to be the major overflow metabolite of LC300 [22] when grown on glucose, which is also predicted by *iGEL603* (Figure 3). When constraining the acetic acid production rate to the rate measured *in vivo*, *iGEL601* predicts lactic acid as an additional byproduct. Since this metabolite is one of the main byproducts of *G. stearothermophilus*, the closest relative of LC300, this prediction is considered reasonable. Furthermore, lactic acid production would also fill the gaps in the carbon and redox balances in earlier published data [21, 22]. This prediction is further supported by the fact that the LC300 genome contains a putative gene for lactate dehydrogenase [20]. Taken together, we thus consider it likely that lactic acid is a previously unidentified major byproduct of the LC300 metabolism, explaining at least some of the missing carbon and degree of reduction in previously reported data.

A number of assumptions have been made when constructing this genome scale model. For example, we chose teichuronic acid as the major part of the cell wall rather than teichoic acid as is the case for *B. subtilis*. The reason for this choice is the lack of an identified glycerol-3-phosphate cytidylyltransferase in LC300, an enzyme necessary for the synthesis of the teichoic acid backbone. Previous studies have shown that the cell wall of some *G. stearothermophilus* strains lacks teichoic acid [48]. Furthermore, four amino acids (Arg, Cys, His, Trp) were missing from the amino acid composition analysis of LC300 [20]. These amino acids are obviously still necessary for growth and they were included in the protein fraction of the biomass component, with their combined amount corresponding to 1% of the total amino acid content. Branched chain fatty acids were excluded in the biomass composition, despite being shown to be part of the *G. stearothermophilus* fatty acid composition [49]. This simplification should not have a major impact on the growth predictions of the model, but is an important aspect to consider if performing engineering for e.g. fatty acid-derived biofuels. It is also unclear which cofactors are necessary for bacterial growth, but we consider the ones presented by Xavier et al. to be a good estimate based on their consensus with earlier reports [34]. The non-growth associated maintenance and P/O-ratio are additional model uncertainties. These were estimated by aligning model predictions with experimental biomass yields, and further experimental work would be required for more accurate determination of these parameters.

Finally, the potential of LC300 as a biorefinery host was evaluated. This was done by modifying *iGEL601* to allow for production of five industrially relevant biochemicals, followed by flux sampling with biochemical production set as objective while keeping biomass formation at 75% of maximum. With the exception of ethanol production on glucose, the predicted specific productivities were on par with or higher than the highest reported specific productivities for all products on both carbon sources. LC300 expresses rapid substrate uptake rates on both glucose and xylose, and the high specific productivities are a result of the high carbon flow through the central metabolism. Hence, to achieve these high productivities *in vivo*, maintaining high substrate uptake while reducing growth rate is key, which may prove challenging *in vivo*. Nevertheless, the predicted productivities presented here gives an indication of the specific productivities achievable through metabolic engineering of LC300. Furthermore, as previously predicted, the production of fumarate and succinate are optimally performed anaerobically. At this point, LC300 is unable to grow in the absence of oxygen [20], and it is likely that substantial metabolic engineering efforts would be required to support anaerobic growth.

The central carbon metabolism reaction fluxes of the modified versions of *iGEL601* were analysed to identify potential metabolic engineering strategies required for achieving the predicted production rates *in vivo*. For 3HB production, flux analysis showed that upregulation of the pentose phosphate pathway combined with downregulation of the citric acid cycle and deletion of overflow metabolite production could result industrially relevant productivity. These predictions are in line with previously reported engineering efforts. For example, increased NADPH availability by overexpression of the oxidative pentose phosphate pathway has been shown to increase 3HB productivity and yield in *E. coli* [39]. The 3HB production pathway includes the reduction of acetoacetyl-CoA to (R)-3-hydroxybutanoyl-CoA using NADPH, and this reaction is further favored by an increase in available NADPH. Furthermore, downregulation of citrate synthase, the first enzyme of the citric acid cycle, has been shown to increase the specific productivity and yield of acetyl-CoA derived products [50, 51].

## 5. Conclusions

In conclusion, we have constructed a genome scale model of the thermophilic bacterium *Geobacillus* LC300, with reaction reversibilities following thermodynamics at the optimum growth temperature. The model follows the LC300 metabolism closely and allows for reliable predictions of growth, metabolite consumption and production corresponding to previously published experimental data. In addition, previously unidentified byproducts were predicted in the LC300 metabolism, closing the carbon and degree of reduction balances. Furthermore, *iGEL601* predicts productivities on par with or higher than previously reported values for five industrially relevant biochemicals. Finally, when analysed for metabolic engineering targets, the model suggests strategies to improve production of 3HB proven experimentally in mesophilic hosts. It is our hope that this model can be used as a tool to further characterize the metabolism and accelerate the metabolic engineering efforts of the *Geobacillus* genus, and shorten the route for the implementation of *Geobacilli* as biorefinery hosts.

## Acknowledgements

The authors thank Dr. Gustav Sjöberg for valuable discussions and advice on the modeling and software development.

This research did not receive any specific grant from funding agencies in the public, commercial, or not-forprofit sectors.

## Author cobtributions

Emil Ljungqvist: Methodology, Software, Data curation, Formal analysis, Visualization, Writing - Original draft; Martin Gustavsson: Conceptualization, Software, Supervision, Writing - review and editing

## Supplementary data

Supplementary file 1.xlxs: Supplementary Tables 1-8.

GitHub repository: The model iGEL601, all modelling scripts, and the resulting data are available as a github repository at https://github.com/margu2/iGEL601/tree/FirstSubmission

